# AI-Generated Hallmarks of Aging and Cancer: A Computational Approach Using Causal Emergence and Dependency Networks

**DOI:** 10.1101/2024.08.28.610200

**Authors:** Jianghui Xiong

**Author notes:** Correspondence: Prof. Jianghui Xiong ( OR).

## Abstract

This study introduces “hallmarks engineering,” a computational approach to generate quantifiable hallmarks of aging and cancer. We evaluated these hallmarks using genome-wide DNA methylation data from ten age-related diseases. Causal emergence analysis revealed that hallmark-level features show stronger disease associations than individual genes, with improvements up to 9.7 orders of magnitude. Hallmark-based models achieved comparable predictive performance with fewer predictors compared to regular pathway-based models. Dependency network analysis uncovered regulatory networks with power-law distributions and identified top-level “super-regulators” such as genomic stability. Notably, the inclusion of neurodegenerative and cancer hallmarks enhanced representation for their respective disease categories. Our findings suggest that top-down modeling using computationally generated hallmarks may reveal common mechanisms across multiple diseases, offering a promising approach for modeling multimorbidity.

## Introduction

A major challenge in biomedicine is synthesizing information from diverse, large-scale datasets to represent the causal mechanisms underlying multiple diseases. This challenge is especially relevant when addressing multimorbidity—the simultaneous presence of two or more medical conditions. Multimorbidity poses significant obstacles for healthcare systems and pharmaceutical research [1]. The current approach of treating diseases in isolation often falls short for elderly patients and can lead to adverse effects due to polypharmacy. There’s an urgent need to develop innovative strategies targeting the fundamental processes underlying age-related multimorbidity, rather than focusing on individual diseases separately. It’s crucial that we shift from the traditional “one drug/one disease” paradigm towards a more holistic, integrated approach [2].

A series of “Hallmarks” papers has profoundly influenced the scientific community. These include the Hallmarks of Cancer [3], Hallmarks of Aging [4], and Hallmarks of Health [5], among others. These publications have gained prominence for their ability to provide a concise, high-level understanding of complex disease-related biological processes. By distilling intricate biological phenomena into fundamental principles, they present a conceptual framework that enhances the accessibility and comprehensibility of the subject matter.

In this paper, we aim to use machine learning tools to generate a quantitative form of hallmarks and use these hallmarks to create a concise representation of multiple age-related diseases. Our computational framework focuses on three questions:

1. How can we preserve information and predictive power for diseases when condensing micro-features into macro-features?
2. Is it feasible to generate a highly simplified representation and predictive model for diseases?
3. From the perspective of disease intervention and drug development, how can we identify critical regulators underlying disease symptoms and pathway alterations, and pinpoint common regulatory variables across multiple diseases?

To address these questions, our computational framework incorporates three key components: causal emergence analysis, parsimony predictive modeling, and dependency network analysis. This study evaluates the potential of AI-generated hallmarks to enhance existing pathways and ontologies in biomedical research infrastructure. We examine their efficacy in identifying superior disease biomarkers and network nodes capable of remodeling diverse pathways, potentially altering disease phenotypes. Moreover, these AI-generated hallmarks may offer a high-level modeling approach to uncover common underlying causal mechanisms across multiple age-related diseases, presenting a novel strategy to tackle the complexities of multimorbidity.

## Results

### AI-Generated Hallmarks and Their Evaluation Framework

Hallmarks represent a higher-level synthesis and refinement of regulatory pathways. In contrast to conventional pathways such as gene ontology terms and signaling pathways, hallmarks function as macro-level features. We developed representative gene sets for hallmarks by consolidating thematically related pathways (**Fig. 1A**). Our hypothesis posits that alterations in these hallmarks constitute the fundamental mechanisms underlying diseases. As macro-level constructs, hallmarks more effectively capture the shared mechanisms across multiple diseases compared to standard pathways. This approach enables the use of a select few hallmarks to elucidate the core mechanisms driving multiple diseases and their associated symptoms, thereby playing a pivotal role in multimorbidity analysis (**Fig. 1A**).

**Figure 1:**
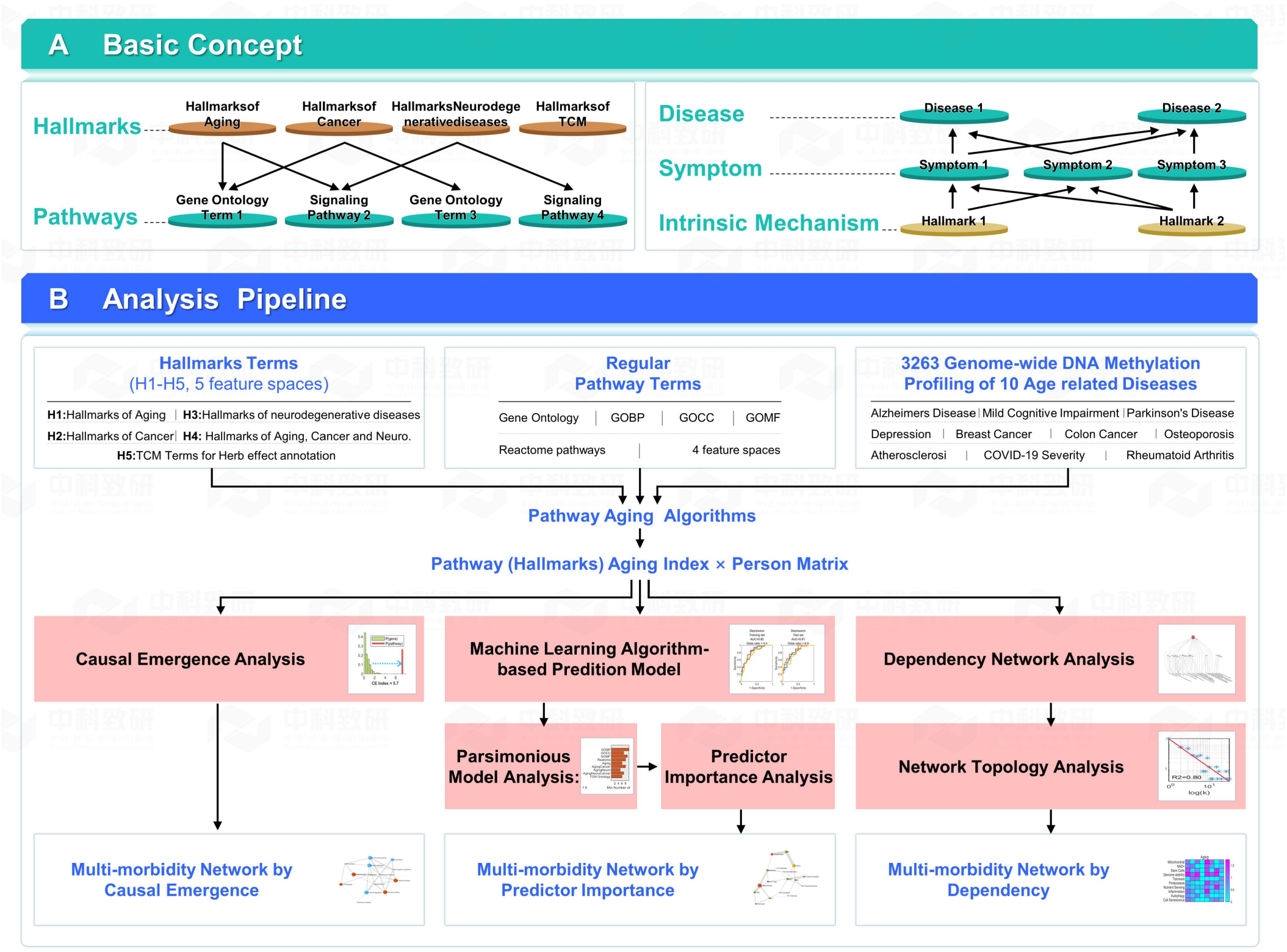
Computational Framework for AI-Generated Hallmarks: Design, Evaluation, and Practical Application.

Our machine learning-based framework for hallmark generation is illustrated in **Figure 1B**. The conceptual foundation for our hallmark terms draws primarily from three influential publications: Hallmarks of Aging [4], Hallmarks of Cancer [3], and Hallmarks of Neurodegenerative Diseases [6]. Our framework incorporates five distinct feature spaces for hallmark analysis:

1. H1 (Aging): Comprises 16 terms, including nine primary categories (e.g., Genome Stability, Proteostasis) and seven secondary categories (e.g., NAD+, immune-related terms).
2. H2 (AgingCancer): Expands on H1 with nine additional terms related to cancer processes, totaling 25 terms.
3. H3 (AgingNeuro): Augments H1 with six neuro-specific terms, resulting in 22 terms.
4. H4 (AgingNeuroCancer): A comprehensive space combining all aforementioned hallmarks, totaling 31 terms.
5. H5 (TCM): Incorporates 39 terms derived from Traditional Chinese Medicine terms (see **Methods**).

Our research employs previously developed pathway aging algorithms [7] to calculate aging indices for both pathways and hallmarks, represented as unique gene sets. We utilize genome-wide DNA methylation profiling data from 10 age-related diseases for our analyses.

The computational framework consists of three primary components:

1. **Causal Emergence Analysis**: This component focuses on preserving and enhancing essential causal relationships and phenotype associations when condensing information from micro-level to macro-level features. It aims to maintain the critical links between biological features and disease outcomes during the compression process.
2. **Parsimony Predictive Modeling**: This approach strives to develop the most efficient and concise model for predicting disease outcomes using a minimal set of predictors or hallmarks. Adhering to the principle of parsimony, we aim to identify a combination of hallmarks that offers optimal predictive accuracy with minimal complexity.
3. **Dependency Network Analysis**: This component investigates the underlying dependencies or supporting factors crucial to disease-defining symptoms. Its objective is to pinpoint key nodes or hallmarks within the biological system that, when targeted, could potentially lead to significant improvements in overall health outcomes. This analysis may uncover promising therapeutic targets or intervention points for addressing aging or disease.

### Causal Emergence: Comparing Hallmarks and Regular Pathway Terms

We developed representative gene sets for each hallmark term by consolidating related pathways. Our aim was to examine whether these hallmark gene sets demonstrate stronger disease associations compared to standard pathway terms. To quantify this enhanced association, we introduced the causal emergence index (CE index). **Figure 2** and **figure 3** presents the results for 8 age-related diseases, with each disease represented by 5 distinct subplots.

**Figure 2:**
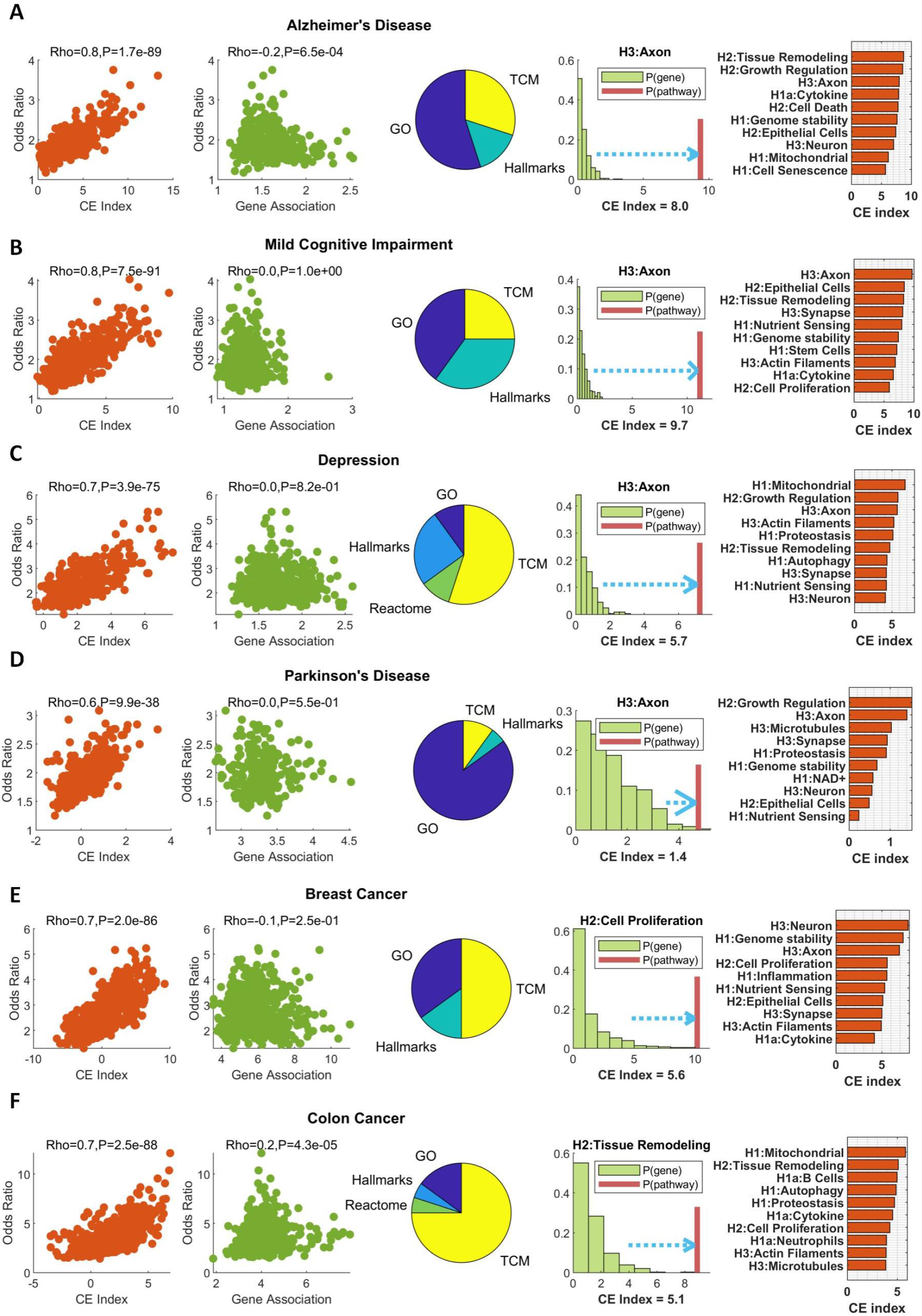
Comprehensive Analysis of Causal Emergence in Hallmarks and Pathways. A–F: Results for six age-related diseases.

**Figure 3:**
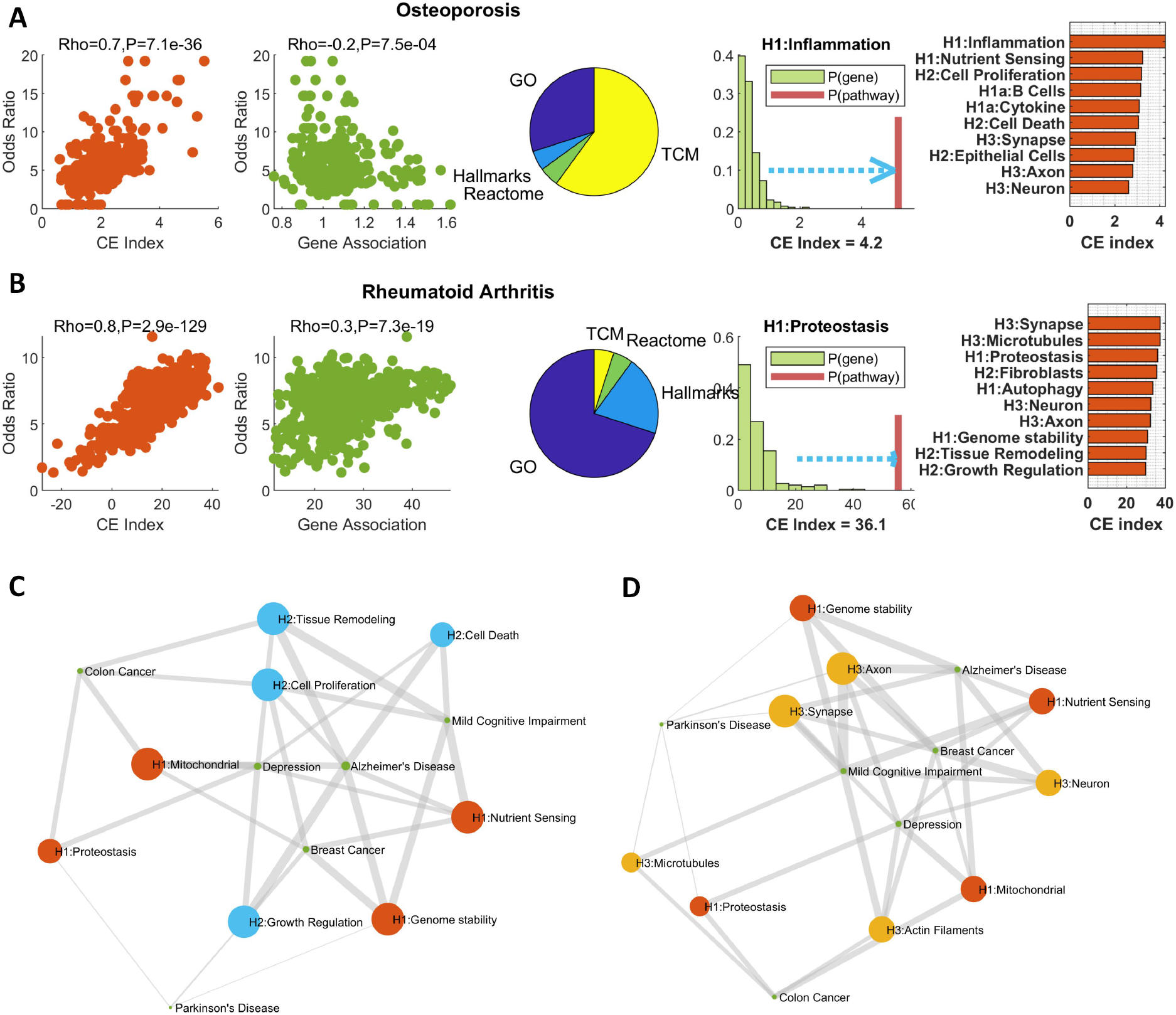
Extended Causal Emergence Analysis and Multimorbidity Insights. A–B: Results for two additional age-related diseases. C: A network visualization of aging and cancer hallmarks associated with multiple diseases. D: A network representation of aging and neurodegenerative disease hallmarks linked to various conditions, highlighting potential multimorbidity mechanisms.

The initial subplot illustrates the relationship between the CE index and odds ratio for each pathway and hallmark. The odds ratio quantifies the association between the aging index of pathways or hallmarks and the disease (methodology detailed in Methods). Across all 8 diseases, we observed a significant positive correlation between the CE index and odds ratio. For example, in Alzheimer’s disease, these variables exhibit a Pearson correlation of ρ = 0.8 (p-value: 1.7E-89). This correlation suggests that pathways more predictive of the disease tend to have higher CE indices.

To address the question of whether a pathway’s strong disease prediction (high odds ratio) is simply due to containing many individually strong predictor genes, we present a second subplot. This green scatter plot explores the relationship between pathway odds ratios and individual gene-disease associations. The gene-disease association is represented by the two-sample t-test p-value of gene variables in the disease group versus healthy control groups. Notably, for most diseases, we found that the pathway odds ratio shows little to no correlation with single gene-disease associations. This observation implies that pathways or hallmarks with high odds ratios likely derive their predictive power from emergent properties rather than individual gene effects.

The third subplot presents pie charts illustrating the distribution of pathways or hallmarks among the top 200 features with the highest CE index. Despite the initial pool of regular pathway terms (GO and Reactome pathways) exceeding 3,000, the hallmarks for aging, cancer, and neurodegenerative diseases comprise only 31 terms. The significant representation of hallmarks within the top 200 CE pathways suggests that these hallmarks contain a higher concentration of high CE features compared to regular pathways.

The fourth subplot depicts a CE principle diagram, while the fifth subplot ranks hallmarks by their CE index. Notably, the hallmark “Axon” (defined by the hallmarks of neurodegenerative disease, H3) consistently ranks in the top three for all neurosystem-related diseases. For example, in Mild Cognitive Impairment, H3:Axon is the top hallmark with the highest CE index of 9.7. In the CE principle diagram, the green bar represents the frequency distribution of gene-disease associations, with the x-axis showing -log10(p), where p is the two-sample t-test p-value of gene DNA methylation values in disease versus control groups. The red line illustrates the association of hallmarks with disease, also as -log10(p), where p is the two-sample t-test p-value of the hallmark aging index in disease versus control groups. The CE index, calculated as per the formula detailed in Methods, represents the difference between the hallmark-disease association score and the 95th quantile of all individual gene-disease association scores. In the case of Mild Cognitive Impairment, H3:Axon enhances the disease association by 9.7 orders of magnitude.

Our analysis indicates that hallmarks with high causal emergence indices exhibit relevance across multiple age-related diseases, highlighting their potential for multimorbidity analysis. By networking the top 10 hallmarks with the highest CE index for each disease, we identified key hallmarks connecting multiple conditions: Genomic Stability (H1) links to Alzheimer’s disease, mild cognitive impairment, Parkinson’s disease, and breast cancer; Tissue Remodeling (H2) connects to Alzheimer’s disease, mild cognitive impairment, depression, and colon cancer (**Fig 3C**); and Axon (H3) relates to Alzheimer’s disease, mild cognitive impairment, Parkinson’s disease, and breast cancer (**Fig 3D**). This cross-disease relevance of hallmarks emphasizes their potential as valuable tools for comprehending and analyzing multimorbidity in age-related diseases.

### Hallmarks Generate Parsimonious Disease Prediction Models

We compared prediction models using hallmarks (macro-level features) with those using regular pathway aging indices. Nine distinct feature spaces were created: three sets of hallmarks (aging, cancer, and neurodegenerative diseases), a combined hallmark set, hallmarks of Traditional Chinese Medicine (TCM), and four regular pathway categories (GOBP, GOCC, GOMF, and Reactome). Each feature space corresponded to a model using only variables from that specific set. For each disease and feature space, we employed LASSO algorithms to generate 10 prediction models by resampling samples 10 times.

Figure 4. illustrates eight disease panels, each containing three subplots. The first subplot in each panel is a scatter plot, with the x-axis representing the mean number of predictors in the 10 resampling-generated models, and the y-axis showing the average predictive performance AUC (area under the ROC curve). Each point denotes a feature space, ranging from standard pathways like GOBP to hallmark series. Generally, a positive correlation exists between the number of predictors and AUC, although increasing variables does not always result in a continuous AUC increase.

**Figure 4:**
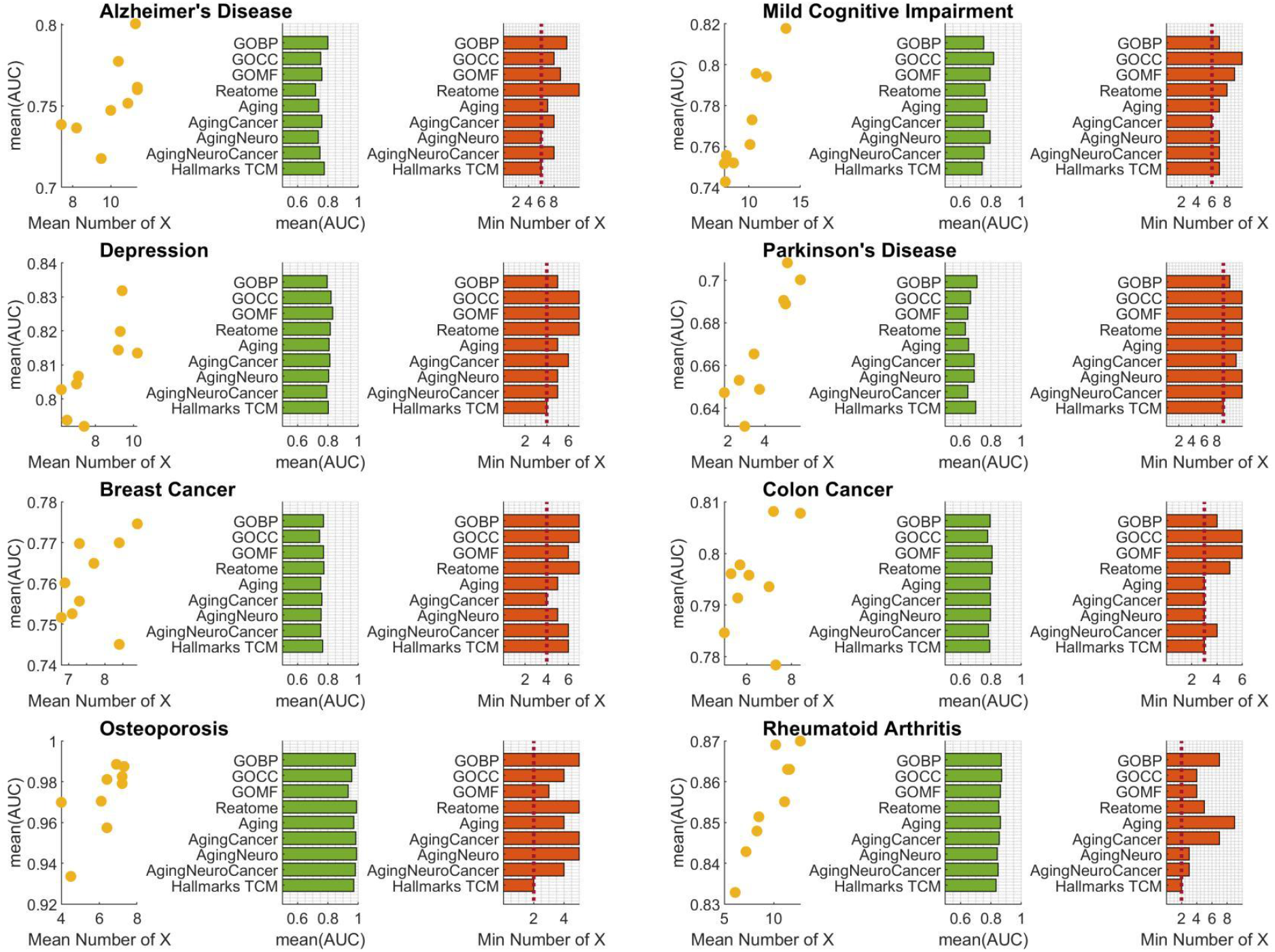
Comparative Analysis of Hallmark and Standard Pathway Feature Spaces for Developing Concise Disease Prediction Models.

The second subplot, a green bar chart, displays the average AUC for the nine feature spaces. Hallmark series feature spaces typically demonstrate AUC levels comparable to standard pathways. To identify which feature space utilizes fewer predictive variables while maintaining similar prediction accuracy, we established a uniform AUC cutoff of 0.75. We then selected models from each feature space exceeding this cutoff and examined the minimum number of predictive variables employed in these models.

The third subplot reveals intriguing patterns across diseases. For Alzheimer’s disease, both AgingNeuro and TCM hallmarks achieve the minimum of six predictive variables. The AgingNeuro feature space, encompassing all aging hallmarks plus several neuron-related ones like H3:Axon, outperforms the aging hallmarks alone. This indicates that neurodegenerative disease hallmarks provide unique predictive variables for neuropsychiatric diseases, enhancing prediction accuracy. Similarly, for breast cancer, AgingCancer reaches the lowest value of four, surpassing the aging hallmarks alone, suggesting that cancer hallmarks offer distinct predictive capabilities. For colon cancer, AgingCancer hallmarks also share the top position, further emphasizing the synergy between aging and disease-specific hallmarks in predictive modeling.

Notably, TCM hallmarks achieve the minimum value in six out of eight diseases studied: Alzheimer’s disease, Depression, Parkinson’s disease, colon cancer, osteoporosis, and Rheumatoid arthritis. This suggests that TCM hallmarks possess distinct advantages in generating simplified predictive models across various diseases.

We utilized two-thirds of the disease dataset for training the prediction models, with the AUC values in **Figure 4** representing training set performance. Test set AUC analysis yielded results consistent with those presented in **Figure 4**. Additionally, when applying a different AUC cutoff (e.g., 0.70) to generate the minimum number of predictors for each disease, the results remained similar to those presented in **Figure 4** (**Supplementary Figure 1**).

### Key Hallmarks Emerge as Predictive Indicators Across Multiple Age-Related Conditions

To enhance model interpretability, we employed the Lasso algorithm, a linear model renowned for its transparency in machine learning. We utilized multiple resampling techniques to generate diverse model iterations, then identified key predictive features using a novel metric: the predictor importance score (detailed in **Methods**).

Our analysis compared two feature spaces: one utilizing classic aging hallmarks, and another incorporating cancer and neurodegenerative disease hallmarks. **Figure 5A** illustrates this comparison, presenting variable importance rankings for both feature spaces across various disease panels.

**Figure 5:**
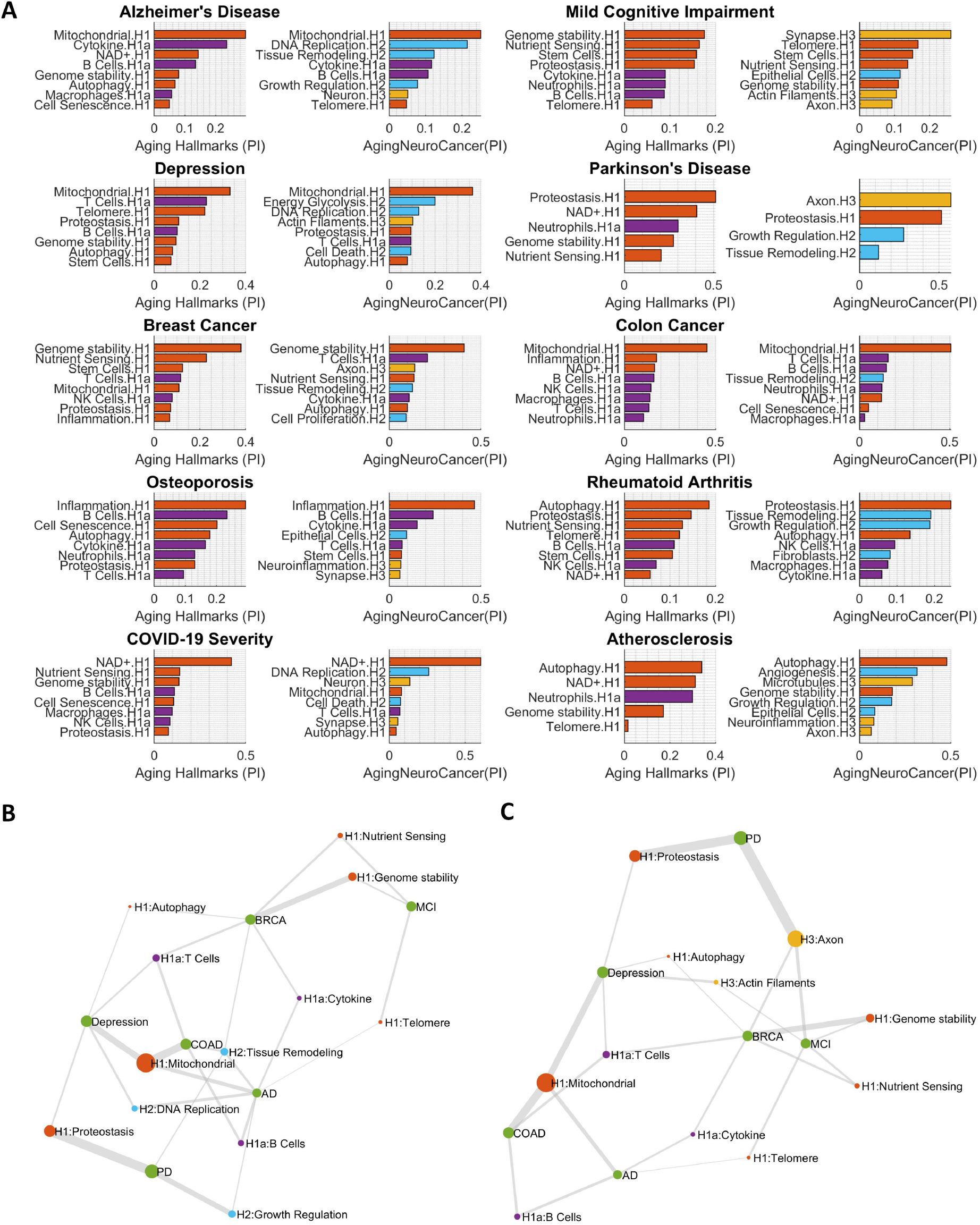
Predictor Importance Analysis. **A**.Comparison of important predictors when adding hallmarks of cancer and neurodegeneration to the basic aging hallmarks.**B**.Multimorbidity analysis on H1 and H2, showing how the combination of important aging and cancer hallmarks connects to multiple age-related diseases.**C**.Multimorbidity analysis on H1 and H3.

In the case of Alzheimer’s disease, mitochondrial factors ranked highest, followed by cytokine and NAD+. The expanded model, which included cancer and neurodegenerative hallmarks, introduced three new predictors from H2 hallmarks: DNA replication, tissue modeling, and growth regulation. These additions underscore the integral role of these processes in normal physiology, extending beyond cancer development. Notably, a Neuron.H3 hallmark emerged as a key predictor in Alzheimer’s disease, while mild cognitive impairment models incorporated additional neural hallmarks such as Synapse, Actin Filaments, and Axon.

For cancer-related conditions, the models highlighted two H2 hallmarks: tissue remodeling and cell proliferation, as significant predictors in breast cancer. Tissue remodeling also demonstrated prominence in colon cancer predictions.

In summary, while neurodegenerative and cancer-specific hallmarks enhance predictive value for their respective disease categories, aging hallmarks consistently maintain their significance as key predictors across various conditions. Notably, mitochondrial function emerges as the top predictive variable for Alzheimer’s disease, depression, and colon cancer, indicating its widespread relevance as a common indicator of multiple age-related diseases. Other hallmarks retaining their prominence include genomic stability for breast cancer, inflammation for osteoporosis, NAD+ metabolism for COVID-19 severity, and autophagy for atherosclerosis.

To visualize these relationships, we constructed networks linking diseases with their most important hallmark predictors. **Figure 5B** depicts the network incorporating aging and cancer hallmarks, with node sizes reflecting cumulative predictor importance across diseases. H1: Mitochondrial stands out as the most prominent node, connecting to multiple conditions. The cancer-specific hallmark H2: Tissue Remodeling demonstrates a unique role by linking breast cancer, colon cancer, Alzheimer’s disease, and Parkinson’s disease. **Figure 5C** illustrates a similar network focused on aging and neurodegenerative hallmarks, where H3: Axon emerges as a significant predictor for Parkinson’s disease, mild cognitive impairment, and breast cancer. These network analyses, derived from machine learning prediction models, offer valuable insights into common indicators across multiple diseases, potentially advancing our understanding of multimorbidity in age-related conditions.

### Dependency Networks Exhibit Power Law Distributions

Intricate interdependencies within biological systems play a crucial role in identifying effective drug targets. These systems comprise complex networks of pathways and processes that influence one another. By understanding these dependencies, researchers can pinpoint key regulatory nodes that, when targeted, may exert broad effects on disease processes. The brain-heart relationship serves as an illustrative example: the brain’s high metabolic demands necessitate a constant supply of oxygen and nutrients, provided by blood circulation driven by cardiac function. Consequently, any impairment in heart function can rapidly compromise the brain’s essential resources, underscoring the critical interdependence of these systems. This interconnectedness highlights the importance of considering systemic effects when developing therapeutic strategies.

In our research, we examine the correlation between the aging index of pathway A and disease manifestation. We posit that if the aging index of pathway B influences the strength of this correlation, pathway A’s disease association is dependent on pathway B. For the purpose of identifying therapeutic targets, we prioritize pathway B. Should a pathway B affect multiple pathway A-disease correlations, it emerges as a promising candidate for therapeutic intervention. Consequently, our network topological analysis emphasizes out-degree analysis, as it effectively captures the impact of pathway B—these potential “super regulators.”

For each disease under study, we develop three distinct versions of dependency networks utilizing the following feature spaces: (1) Pathway network: Employing pathway aging indices of standard pathways. (2) Hallmarks aging network: Utilizing a combination of aging indices from standard pathways and hallmarks of aging, cancer, and neurodegeneration. (3) Hallmarks TCM network: Incorporating aging indices from standard pathways and Traditional Chinese Medicine (TCM) hallmarks.

Our dependency network analysis examines all possible two-variable combinations and calculates their dependency index. Network topological analysis results are presented in **Figure 6**. Each disease panel comprises four subplots, with the first displaying log-log plots for the out-degree of dependency networks derived from regular pathway feature spaces. These power-law distribution analyses use log10(k) for the x-axis (k being the out-degree) and log10(Pk) for the y-axis (probability of out-degree). All eight diseases exhibit good linear fit in log-log plots, indicating power-law distribution of out-degree. Colon cancer shows the best linear fit (R-squared: 0.86), followed by Alzheimer’s disease (R-squared: 0.82). Hallmarks aging and TCM feature spaces also demonstrate good power-law distribution fits.

**Figure 6:**
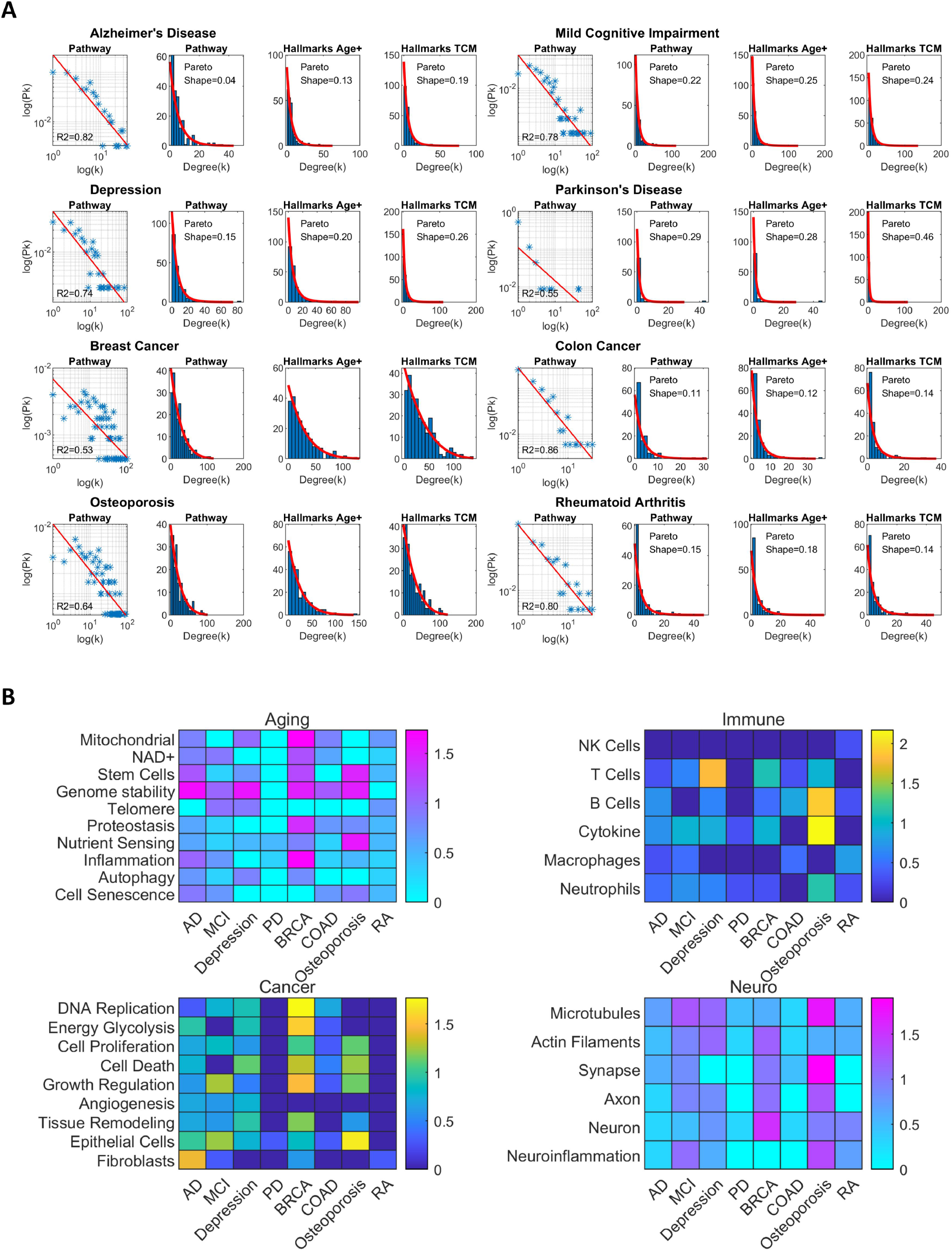
Dependency Network Analysis and Feature Importance. **A**.Degree distribution analysis. **B**.Key regulator nodes in the network across multiple diseases.

To identify potential therapeutic targets, we apply a Generalized Pareto Distribution to the out-degree distribution, a method commonly employed for modeling extreme values in networks and identifying super nodes. The shape parameter, a key output, indicates the impact of highly connected nodes when positive. We compare this parameter across three feature spaces: regular pathways, aging hallmarks, and Traditional Chinese Medicine (TCM) hallmarks.

Six of the eight studied age-related diseases show positive shape parameters. The aging hallmarks feature space consistently demonstrates larger shape parameters compared to regular pathways, indicating enhanced impact of super nodes when hallmark terms are incorporated. For instance, in Alzheimer’s disease, the shape parameter increases from 0.03 (pathway feature space) to 0.13 (aging hallmarks feature space), while in depression, it rises from 0.15 to 0.20.

Notably, the TCM hallmarks feature space often exhibits even larger shape parameters. In Parkinson’s disease, for example, the parameter increases from 0.28 (aging hallmarks) to 0.46 (TCM hallmarks). Similar increases are observed in Alzheimer’s disease (0.13 to 0.19) and depression (0.20 to 0.26).

The above results present the out-degree distribution (**Figure 6**). A similar analysis conducted on the in-degree distribution also conforms to a power-law distribution for some diseases, as shown in **Supplementary Figure 2**.

### Dependency Networks Reveal Critical Regulatory Nodes with Therapeutic Potential

To offer a comprehensive overview of the highly connected nodes in the dependency network, we generated a heatmap illustrating the outdegree of each hallmark term across various diseases. **Figure 6B** presents these findings, categorized into four distinct panels: classical aging hallmarks, cancer hallmarks, neurodegenerative disease hallmarks, and immune function. This segmentation facilitates a more refined analysis of hallmark interactions across different disease categories.

Within the classical aging hallmarks, genomic stability consistently appears as a primary regulatory node. Additional significant terms include stem cells and mitochondrial-associated hallmarks, underscoring their importance in age-related processes.

In the cancer hallmarks category, growth regulation and cell death mechanisms demonstrate prevalence across multiple diseases, with particular prominence in breast cancer. Notably, within the neurodegenerative disease hallmarks, osteoporosis displays high outdegree node terms, specifically in microtubules and synapses. Interestingly, breast cancer also exhibits a highlighted neuronal term, suggesting potential cross-category interactions.

The immune function panel reveals that osteoporosis is characterized by high outdegree in B cells and cytokine-related terms. Concurrently, T cells show significant impact in depression, indicating the diverse roles of immune components across different age-related conditions.

Our analysis revealed the presence of highly influential nodes within the dependency network for each disease. To visualize these relationships, we created partial network diagrams, as shown in **Figures 7** and **8**. Given the complexity of the networks, we prioritized nodes based on their total degree (the sum of outdegree and indegree), focusing on the most significant components. The size of each node in the diagrams corresponds to its total degree, providing a visual representation of its importance. We employed a layered layout to effectively illustrate the hierarchical and regulatory relationships within the networks.

**Figure 7:**
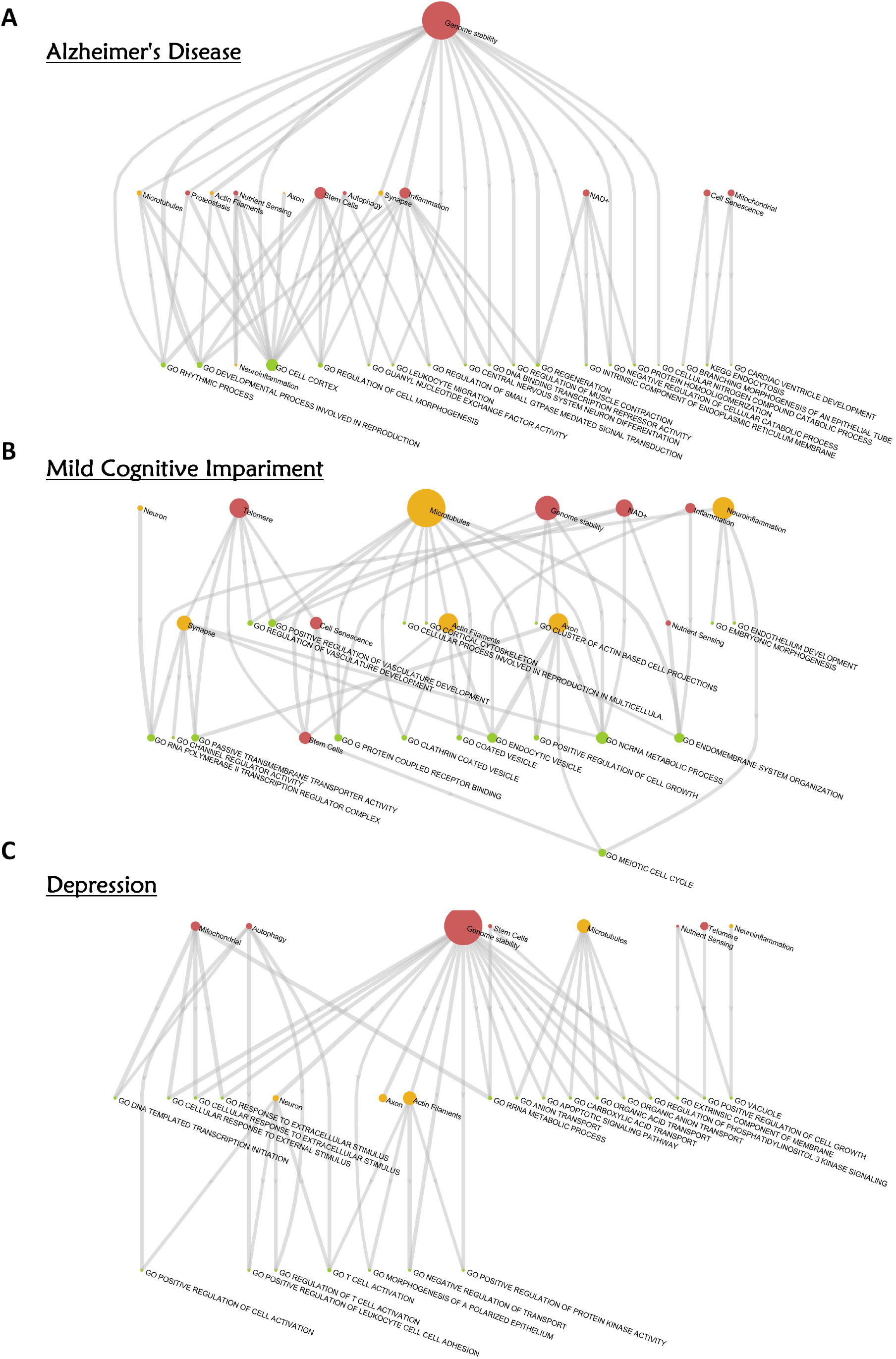
Hallmark Dependency Networks for Neurodegenerative Diseases. **A**.Alzheimer’s disease. **B**.Mild cognitive impairment. **C**.Major depressive disorder.

**Figure 8:**
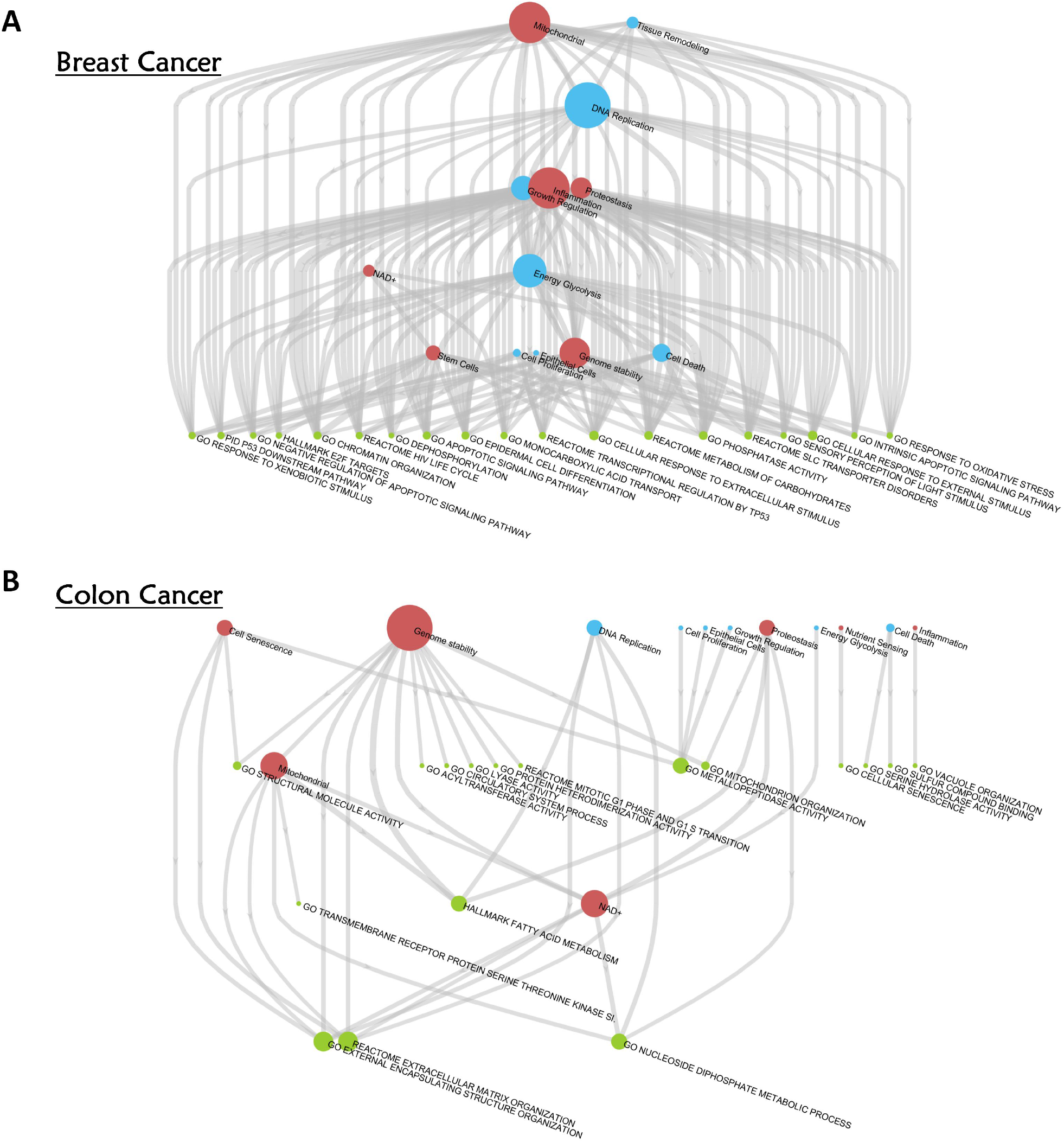
Hallmark Dependency Networks for Cancer-Related Diseases. **A**.Breast cancer. **B**.Colon cancer.

Figure 7. presents the dependency networks for three neurological disorders. We utilized a color-coding system to differentiate between various types of hallmarks: red for classical aging hallmarks, orange for neurodegenerative disease hallmarks, and green for regular pathway terms.

In the case of Alzheimer’s disease (**Figure 7A**), the network exhibits a clear three-tiered structure. At the apex, we find a single node representing genomic stability, which exerts significant influence over nodes in the second and third tiers. The second tier features stem cells and inflammation as key players. Within the third tier, the ‘GO cell cortex’ node stands out due to its larger size, indicating a high number of incoming connections and suggesting that it is subject to regulation by numerous other factors in the network.

The dependency network for mild cognitive impairment reveals a more sophisticated regulatory structure. While genomic stability maintains its crucial role, other nodes, notably microtubules, emerge as comparably significant regulators. This increased variety of key regulators potentially reflects the complex nature of early cognitive decline, suggesting multiple pathways or origins for the condition. Although this study does not primarily aim to identify specific disease mechanisms, it demonstrates the efficacy of dependency network analysis in unraveling the intricacies of disease progression.

In major depressive disorder, genomic stability remains a critical factor. Additional top-tier regulators within classical aging hallmarks encompass mitochondrial function, autophagy, telomeres, and nutrient sensing.

**Figures 8** present the findings for two cancer types, with cancer hallmark terms highlighted in light blue. For breast cancer (**Figure 8A**), mitochondrial function emerges as the most prominent classical aging hallmark. Within cancer hallmarks, DNA replication and energy glycolysis stand out as key nodes. In colon cancer (**Figure 8B**), genomic stability takes the lead as the primary regulator.

Consistent with the original rationale for introducing hallmarks of cancer and aging, our analysis indicates that a select group of key players may elucidate mechanisms in aging-related diseases. The hierarchical structure observed in dependency networks suggests that future research should prioritize interventions targeting top-tier regulators, potentially offering a strategic approach for identifying therapeutic targets.

## Discussion

Recent advancements in identifying hallmarks of aging and cancer represent a significant breakthrough in biomedicine, providing a unified framework for understanding disease mechanisms, identifying biomarkers, and discovering therapeutic targets. This progress is poised to influence drug discovery, personalized preventive medicine, and age-related interventions. Our findings demonstrate the feasibility of employing machine learning techniques to generate data-driven hallmark definitions. This approach paves the way for a novel field of study, termed “hallmarks engineering,” which focuses on developing, assessing, and refining the hallmarks knowledge framework.

The issue of multimorbidity is of particular importance in this context. Current AI approaches to drug discovery often adhere to a “one disease, one drug” paradigm, which may not adequately address the complexities of multiple coexisting conditions. There is a pressing need for more comprehensive, patient-centered models that can analyze multimorbidity, identify relevant predictors, and elucidate underlying pathogenic mechanisms [8]. Our proposed hallmarks engineering method offers a promising solution by providing a macro-level, top-down perspective on biomarkers, regulatory networks, and potential therapeutic targets, thereby facilitating a more holistic approach to addressing multimorbidity.

From a methodological perspective, our previously introduced pathway aging algorithm allows for the assessment of intrinsic capabilities across a wide range, including protein complexes, organelles, organs, and inter-organ systems. We introduce the concept of “capomics”—the study of intrinsic capabilities as defined by pathway or hallmark aging in an omics-like manner. This approach, involving comprehensive analysis of pathway aging for individual patients, shows great promise for personalized health interventions and disease prevention strategies. As demonstrated by the dependency network analysis in **Figures 7** and **8**, a small number of highly influential regulators can significantly impact a wide range of network nodes. Identifying and characterizing these key regulatory elements represents a crucial focus for future research in hallmarks engineering and capomics analysis.

A notable aspect of this study is the comparative analysis between hallmarks of aging and Traditional Chinese Medicine (TCM) hallmarks. The TCM hallmarks demonstrate high causal emergence and rank prominently in predictor importance analyses (**Supplementary Figure 3**), indicating their robust predictive capabilities. In the dependency network analysis, TCM hallmarks exhibit higher Pareto shape parameters compared to other hallmark categories, suggesting the presence of more influential regulatory elements within the TCM framework. The analysis identifies several TCM hallmarks, represented by nodes with high out-degree, that consistently appear across multiple disease networks. This finding, as illustrated in **Supplementary Figure 4**, highlights the potential importance of these TCM concepts in comprehending and addressing various age-related conditions. The construction of a hallmark-exclusive dependency network provides a macro-level perspective on the regulatory interactions between aging and TCM hallmarks (**Supplementary Figures 5** and **6**). TCM hallmarks offer a succinct representation of disease mechanisms and herbal effects, warranting further investigation into their specific merits and potential synergies with pathway-aggregated hallmarks derived from gene ontology and signaling pathways. Elucidating the effectiveness of TCM hallmarks could yield valuable insights for enhancing the overall hallmarks framework, thereby addressing key objectives in both hallmarks engineering and capomics analysis.

This study has certain limitations that should be acknowledged. Firstly, the sample size for each disease is relatively limited. Ideally, multiple cohort datasets per disease would be available to conduct independent validation of the prediction models. Given the constraints in data availability, we opted for a resampling approach and employed the LASSO machine learning algorithm to train a series of prediction models. This methodology enabled us to identify statistical patterns in predictor usage, forming the basis for our predictor importance analysis.

Secondly, the scope of age-related diseases included in this study is not exhaustive. While multimorbidity often involves multiple disease clusters, our research primarily focused on several neurological conditions and two types of cancers. Future research efforts aim to expand the range of diseases studied, with a particular emphasis on including cardio-metabolic disorders to provide a more comprehensive analysis of age-related multimorbidity.

## Methods

### Pathway Aging Analysis

This study utilizes DNA methylation and pathway/ontology data consistent with our previous research [7]. We sourced regular pathway terms from The Molecular Signatures Database (MSigDB) [9], accessible at http://www.gsea-msigdb.org/gsea/msigdb/index.jsp. These encompass gene set definitions from Gene Ontology (GO), including Biological Process (GOBP), Molecular Function (GOMF), and Cellular Component (GOCC). REACTOME pathways were employed to represent signaling pathway terms. For the purposes of this study, we use the term “pathway” to encompass both gene ontology terms and signaling pathway terms.

We interrogated 10 age-related diseases using genome-wide DNA methylation profiling of clinical cohorts. The data was downloaded from the GEO database. Unless specified otherwise, these data are derived from whole blood samples, using the Illumina 450k/850k microarray platform. The total number of clinical samples is 3263. The data examined in this study encompasses several conditions, including Alzheimer’s Disease [10] (GSE153712, N=632), Mild Cognitive Impairment [10] (GSE153712, N=565), Parkinson’s Disease [11] (GSE111223, N=259, Saliva), Osteoporosis [12](GSE99624, N=48), Breast Cancer (GSE51032, N=659), Colon Cancer (GSE51032, N=590), Atherosclerosis [13] (GSE46394, N=49), Depression [14](GSE113725, N=194), COVID-19 Severity [15](GSE179325, N=473), and Rheumatoid Arthritis [16] (N=689).

Each gene ontology and signaling pathway consists of a specific set of genes. We assign a characteristic value to each gene, such as its DNA methylation level, calculated as the average methylation value of all promoter region CpG sites. Utilizing a comprehensive DNA methylation dataset that includes age information, we determine the correlation between each gene and age. This analysis identifies genes positively (pos genes) and negatively (neg genes) correlated with age. A pathway is considered to exhibit more pronounced aging when pos genes show higher characteristic values and neg genes show lower values. To quantify this, we employ a T-test to compare the characteristic values of pos and neg genes, using the resulting T-score as the pathway’s characteristic value.

The aging of each pathway is determined by the statistical difference between the DNA methylation values of Pos-genes and Neg-genes. We conduct a T-test between these values for each DNA methylation profile. The resulting T-statistics serve as the pathway aging index (PA index). Specifically, **x** represents DNA methylation values of Pos-genes, and **y** represents those of Neg-genes. The two-class t-test of gene scores (DNA methylation scores) between sets **x** and **y** is calculated using the following T-statistic:

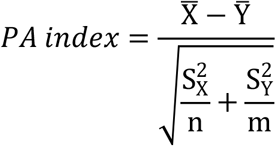

Here, 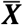 is the mean of gene scores of gene set ***X***, 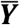 is the mean of gene scores of gene set ***Y***. 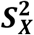 is the variance of gene set X scores, 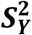 is the variance of gene set ***Y*** scores, ***n*** is the number of genes in set ***X***, and ***m*** is the number of genes in set ***Y***.

This t-score serves as the disease-specific pathway aging index. Through this process, we transform each disease’s methylation profile into a new matrix—the PA-index by samples matrix—where each row represents a pathway aging index.

### Aggregating Pathways into Hallmark Gene Sets

We define gene sets for established hallmarks of aging and cancer by consolidating groups of standard pathways. These pathway definitions are sourced from gene ontology or signaling pathways in the GSEA database. For each hallmark, we meticulously select relevant keywords. For instance, to represent “mitochondrial dysfunction” in the aging hallmarks, we employ “mitochondrial” as a search term in the GSEA database to identify all pathways containing this keyword. Pathways annotating 50 or more genes contribute their entire gene set to a comprehensive list. Genes essential to mitochondrial function frequently appear multiple times in this compilation. We subsequently calculate each gene’s frequency of occurrence, rank them in descending order, and select the top 1000 genes as the representative set for that particular hallmark. Given that many of the diseases under analysis are neurodegenerative in nature, we also incorporate the Hallmarks of neurodegenerative diseases into our analysis.

In the realm of Traditional Chinese Medicine (TCM), practitioners utilize a concise set of terms to characterize the functions or effects of herbs on the human body. These terms serve to group similar herbs, and herbs sharing the same term annotation often influence overlapping biological processes at the cellular level. This observation suggests that a specific TCM term could potentially represent a distinct gene set. We applied this concept to generate a hallmark gene set for each TCM term.

To accomplish this, we developed a comprehensive TCM functional annotation database. This database was created by extracting pertinent keywords from the “function” sections of TCM notes found in the Chinese Pharmacopoeia (502 species), the National Collection of Traditional Chinese Medicine (3741 species), and the Chinese Materia Medica (7832 species). From this extensive collection of TCM terms, we carefully selected 39 keywords based on their frequency of occurrence in the annotation database, with a particular emphasis on syndrome descriptions.

For each TCM term, we compiled a list of associated herbs. We then obtained the chemical composition of these herbs using the TCMID database [17]. For each chemical, we retrieved its target gene list from the STITCH database [18], applying a STITCH score threshold of ≥200. This process yielded a target gene list for each TCM term. Genes crucial to a specific TCM term often appeared multiple times in these lists. We calculated each gene’s frequency of occurrence, ranked them from highest to lowest, and selected the top 1000 genes as the representative set for that hallmark.

### Causal Emergence Analysis

Causal emergence, a concept in complexity science, suggests that macro-scale phenomena can enhance causal relationships by reducing noise, resulting in stronger causal links at higher organizational levels [19]. In our study, we consider pathway-level features as macroscale phenomena and individual gene events as microscale features. We further aggregate pathways into hallmarks, representing an even higher macro level. Our aim is to evaluate whether the association of pathway (or hallmark) aging index with phenotype (disease) is enhanced compared to individual genes.

To assess gene-disease associations, we developed a gene-by-person matrix for each disease, with matrix values representing gene DNA methylation levels. We utilized the average methylation value of all promoter region sites as the gene’s representative value. For each gene, we computed the t-test p-value comparing methylation levels between the disease and healthy control groups, using -log10(p) as the association index (**Ag**).

For pathway-disease associations, we employed a similar methodology, substituting gene methylation values with pathway aging index scores. We began with a pathway aging index by persons matrix and used -log10(p) as the pathway-disease association index (**Ap**). Hallmarks-disease associations were calculated analogously, using the hallmarks aging index.

Consequently, each pathway/hallmark has a single pathway-disease association index value, while each pathway encompasses multiple gene-disease association indices. We then defined the causal emergence index (CE index) for each pathway/hallmark to quantify the difference between macro-level (pathway-disease) and micro-level (gene-disease) associations. The CE index is computed as **Ap - Agr**, where Agr represents the 95th percentile of **Ag**.

In our causal emergence analysis, we also examined the relationship between the CE index and odds ratio. We utilized the odds ratio to indicate the association between the pathway or hallmarks aging index and disease. To calculate this, we determined the average pathway aging index for each pathway across all samples, then categorized samples into PA-High and PA-Low groups based on this average. Using Fisher’s exact test, we quantified the difference in disease probability between these groups, yielding an odds ratio.

### Disease Prediction Modeling and Predictor Importance Analysis

Our methodology involves conducting 10 resampling iterations for each disease and feature space, such as hallmarks of aging. For each iteration, we randomly assign 2/3 of the disease dataset samples to construct a predictive model using the LASSO algorithm, while the remaining 1/3 serves as a validation set. This approach yields 900 predictive models across 10 diseases, 9 feature spaces, and 10 models per combination, establishing a comprehensive framework for analysis across various conditions and features.

To assess the relative importance of predictor variables, we calculate a Predictor Importance (PI) index for each variable across the 10 resampling iterations within a specific disease and feature space. The PI index for the i-th predictor variable is quantified as follows:

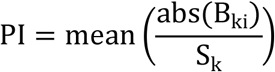

Here, **k** denotes the k-th model (1 to 10), **Bki** is the coefficient of the i-th predictor variable in that model, abs represents the absolute value operation, and **Sk** is the sum of absolute values of all predictor variable coefficients in the k-th model. The mean function computes the average value across all models.

### Dependency Network Analysis

In our study, we examine the relationship between variables and disease phenotypes. We consider a variable Va to be dependent on another variable Vb if Vb influences the correlation between Va and the phenotype. For pathways a and b, we assess this dependency using their aging indices and disease association. Our dependency network analysis follows a two-step process for each disease:

Step 1: Pathway ranking. We calculate the average pathway aging index for each pathway across all samples. Samples are divided into PA-High and PA-Low groups based on this average. Using Fisher’s exact test, we determine the difference in disease probability between groups, obtaining an odds ratio. Pathways are ranked by odds ratio, with the top 200 selected for further analysis.

Step 2: Dependency index calculation. For each pathway pair, we assume their aging indices are Va and Vb. To test whether Va’s correlation with the disease depends on Vb, we first calculate Vb’s average value across all samples. We classify samples above this average as the Vb-High group and those below as the Vb-Low group. Within these groups, we assess the correlation between Va and the disease phenotype using two-sample t-tests. We then calculate the difference between the T-scores of these tests, as shown in the following formula:

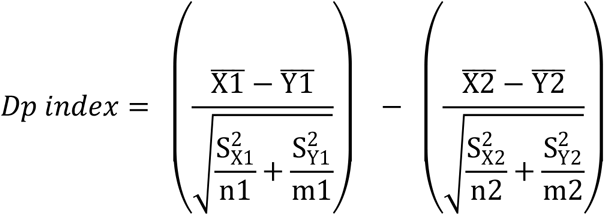

In this formula, the strength of dependency is the difference between two T-scores. The larger this difference, the greater the impact of Vb’s state on the correlation between Va and the disease. The first parenthesis represents the calculation for the Vb-high group, while the second represents the Vb-low group. **X1** is Va (the aging index of pathway a) in the disease group of the Vb-high population, **Y1** is Va in the healthy control group of the Vb-high population. 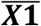 and 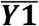 are the means of **X1** and **Y1**, respectively. 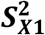 and 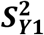 are the variances of **X1** and **Y1. n1** and **m1** are the number of samples in **X1** and **Y1**. Similarly, in the second parenthesis, **X2** is Va in the disease group of the Vb-low population, **Y2** is Va in the healthy control group of the Vb-low population. 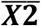 and 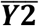 are the means of **X2** and **Y2**. 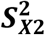 and 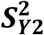are the variances of **X2** and **Y2. n2** and **m2** are the number of samples in **X2** and **Y2**.

In our dependency network analysis, we compute a dependency index for all possible variable pairs. A dependency relationship between two pathways is established when this index surpasses a predetermined threshold (e.g., 3). This relationship is represented as an edge in a directed network.

To analyze the network’s structure, we examine the degree distribution using log-log plots for both out-degree and in-degree. We employ the Generalized Pareto Distribution to model extreme values in the network, which aids in identifying potential therapeutic targets. The shape parameter of this distribution provides insight into the influence of highly connected nodes. By fitting the out-degree and in-degree distributions to this model, we obtain shape parameters that quantify the importance of super nodes within the network architecture.

## Data Availability

All data used in this study are derived from publicly available sources. Detailed access information for these datasets are provided in the Methods section of this paper.

## Acknowledgement

This paper has not received any external funding.

