## Supplemental Figures for "AI-Generated Hallmarks of Aging and Cancer: A Computational Approach Using Causal Emergence and Dependency Networks"

### Supplementary Figures

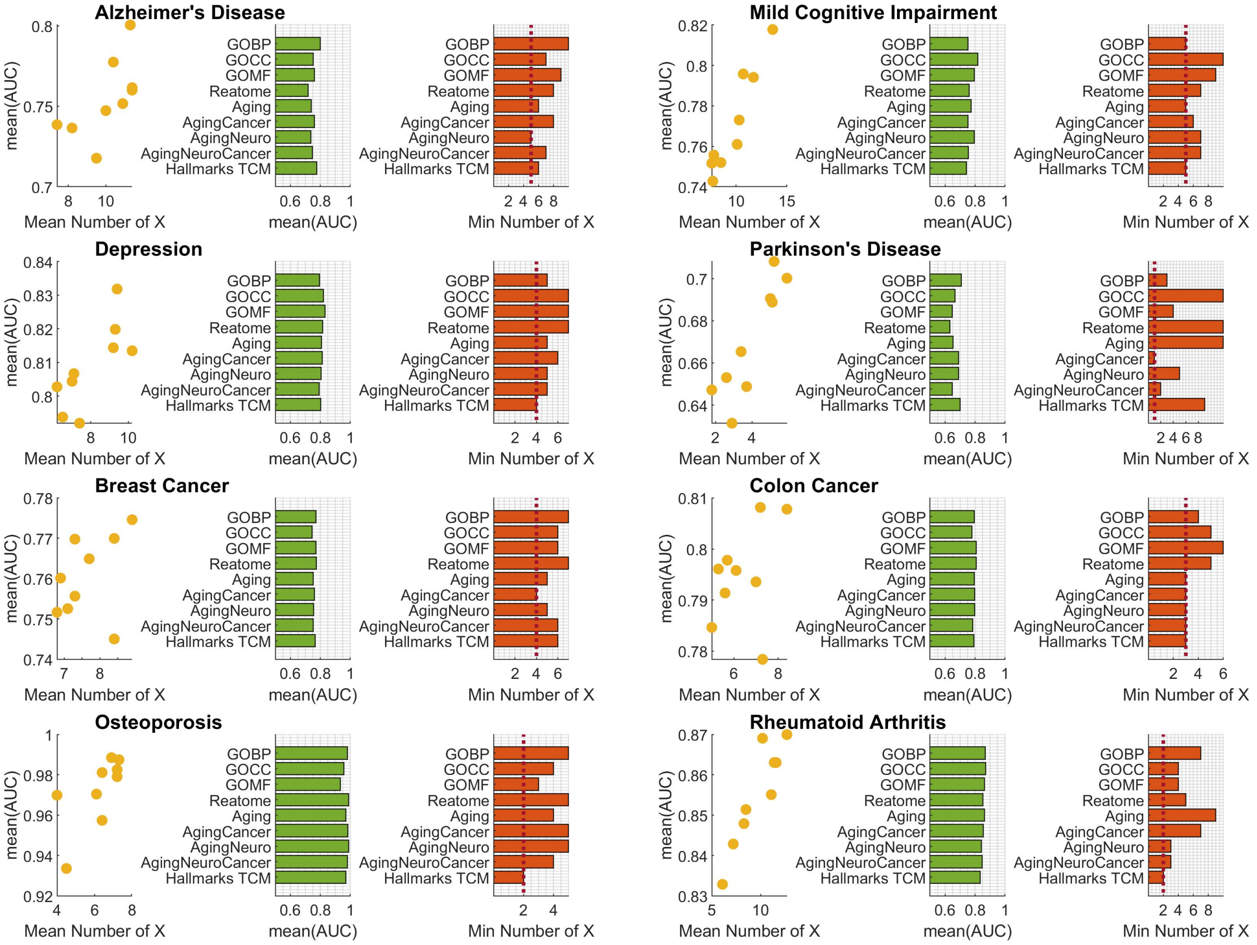


**Supplementary Figure 1: Evaluating Hallmarks and Standard Pathway Feature Spaces for Developing Concise Disease Prediction Models.** This plot is similar to Figure 4 in the main text, with the key difference being the use of an AUC cutoff of 0.70 to generate the minimum number of predictors for each disease.

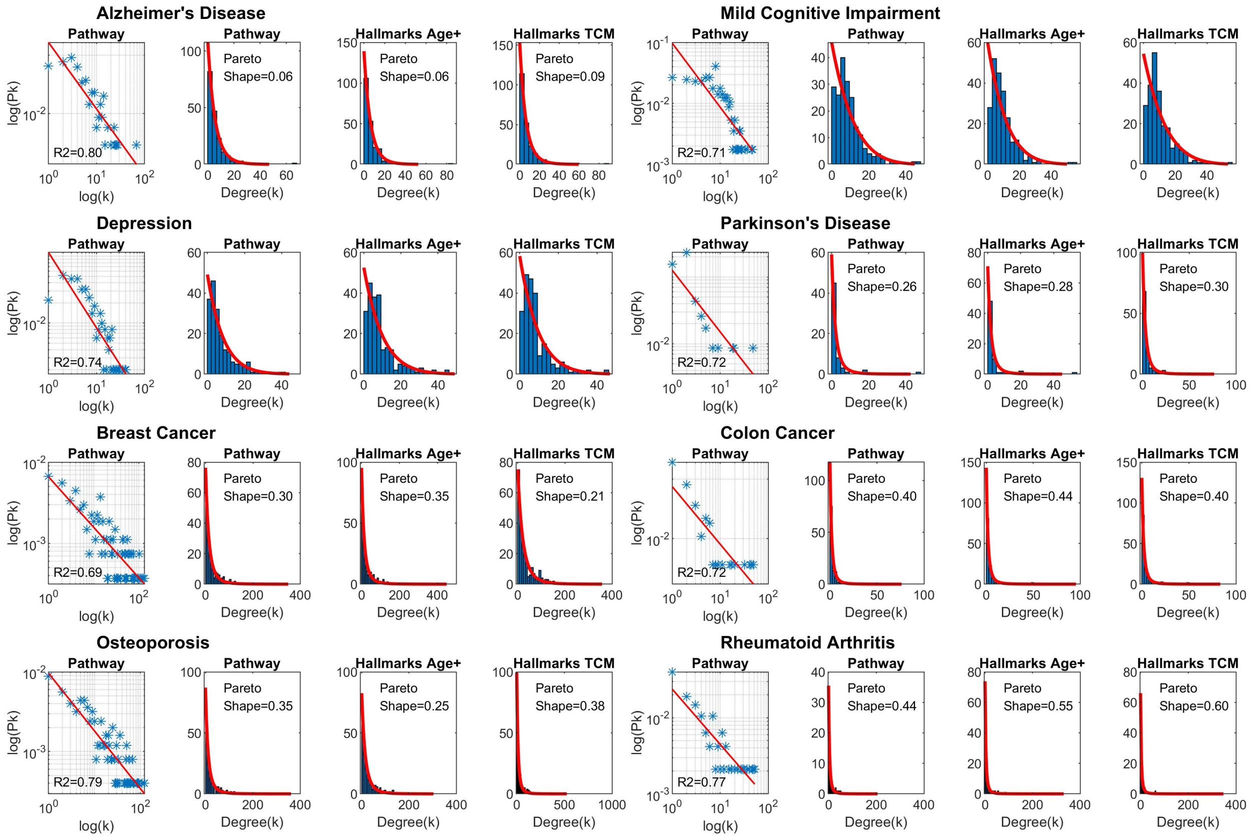


**Supplementary Figure 2: Dependency Network Analysis for In-Degree Distribution.** This plot is similar to Figure 6 in the main text, with the key difference being that Figure 6 analyzes the out-degree distribution to identify key regulators, while this plot examines the in-degree distribution.

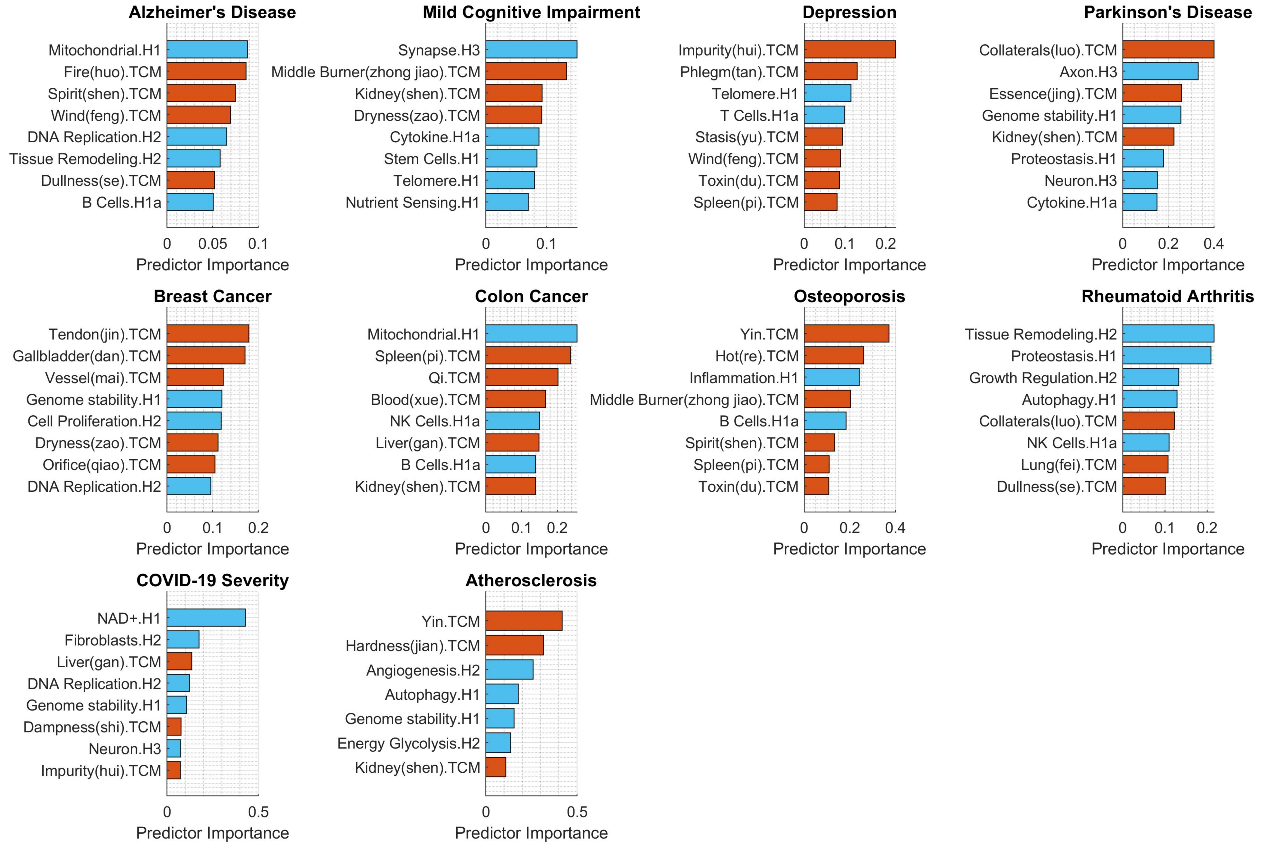


**Supplementary Figure 3: Predictor Importance Analysis Using TCM Hallmarks.** This analysis combines feature spaces of hallmarks from aging, cancer, and neurodegeneration (ageCancerNeuro) with Traditional Chinese Medicine (TCM) hallmarks to create a mixed feature space. We then rank all predictors by their predictor importance index to compare the rankings of TCM hallmarks (shown in red) against ageCancerNeuro hallmarks (displayed in light blue).

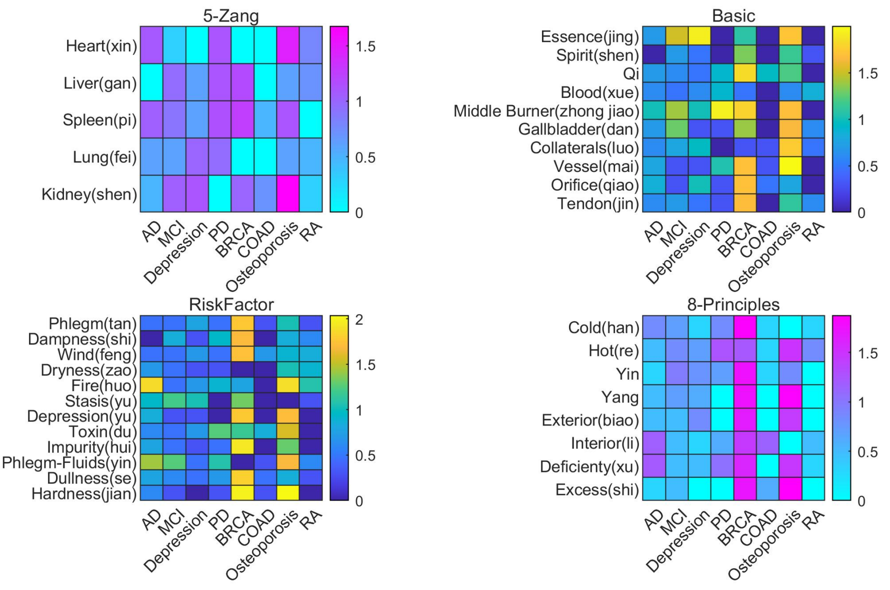


**Supplementary Figure 4: Key regulator nodes as TCM hallmarks in the network across multiple diseases.** Heatmap for investigating the out-degree of TCM hallmark nodes in the dependency networks across multiple diseases.

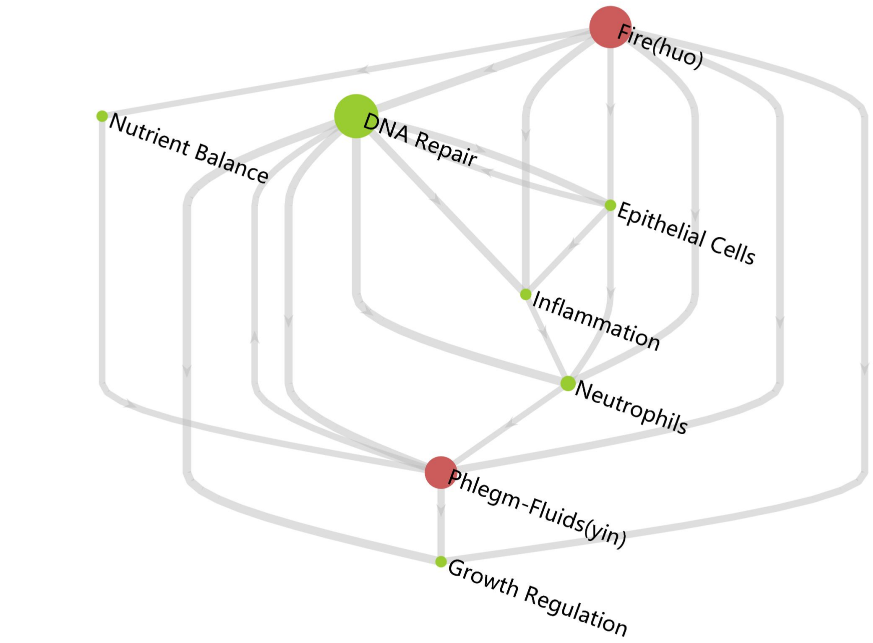


**Supplementary Figure 5: Top-level dependency network between hallmarks of aging and TCM hallmarks for Alzheimer's disease.** In contrast to the main text's dependency network—which included regular pathways and hallmarks of aging—this figure presents a more focused view. It exclusively shows relationships between hallmarks of aging and TCM hallmarks, omitting regular pathway nodes. This approach yields a concise, top-level view of the hallmarks regulation.

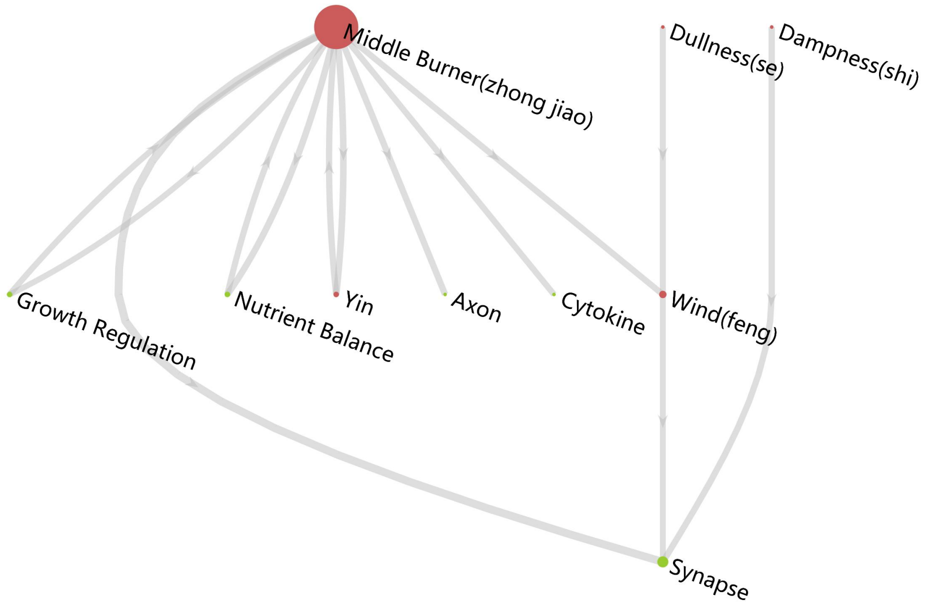


**Supplementary Figure 6: Top-level dependency network between hallmarks of aging and TCM hallmarks for Parkinson's disease.** This plot was generated using similar logic to Supplementary Figure 5. The key difference is that this network focuses on Parkinson's disease, providing a concise, top-level view of the hallmarks regulation for this condition.
